# Automating Morphological Profiling with Generic Deep Convolutional Networks

**DOI:** 10.1101/085118

**Authors:** Nick Pawlowski, Juan C Caicedo, Shantanu Singh, Anne E Carpenter, Amos Storkey

## Abstract

Morphological profiling aims to create signatures of genes, chemicals and diseases from microscopy images. Current approaches use classical computer vision-based segmentation and feature extraction. Deep learning models achieve state-of-the-art performance in many computer vision tasks such as classification and segmentation. We propose to transfer activation features of generic deep convolutional networks to extract features for morphological profiling. Our approach surpasses currently used methods in terms of accuracy and processing speed. Furthermore, it enables fully automated processing of microscopy images without need for single cell identification.

## 1 Introduction

Modern microscopy and automation technologies enable large-scale experiments which can produce millions of cell images per day [7]. This changes the way biologists analyze microscopy images from targeted measurements (‘screening’) to a more wide-ranging feature extraction (‘profiling’) that aims to capture a broad set of measurements representing the cellular phenotypes. Such ‘unbiased’ representations of phenotype enable new analysis based on similarities and differences among chemical or genetic perturbations, which might lead to new insights in functional genomics, drug discovery and target identification [1].

Morphological profiling can be interpreted as a computer vision task to extract relevant features from microscopy images. Many current approaches use specialized software such as CellProfiler [3] and rely on hand-tuned segmentation and feature extraction for each assay [6]. Recent successes in computer vision, however, were driven by learning features directly from data [4], [8], [14], an approach that can be applied to profiling.

In this work, we study the problem of transferring features learned on natural images [11] to obtain unbiased morphological profiles from fluorescence microscopy images of cultured human cells. By obviating the need for single cell identification, the proposed approach offers the following advantages:

- *Speed:* It enables faster profile extraction than the classical pipeline with image segmentation and feature extraction.
- *Autonomous:* It eliminates the need for human input to tweak parameters.
- *Performance:* The extracted profiles achieve a better accuracy than a baseline approach based on handcrafted features.

## 2 Related work

This work lies at the intersection of morphological profiling and deep learning. In morphological profiling, Ljosa et al. [9] compared various profiling algorithms using the same experimental setup, which is also used in the current paper (BBBC021 [2]): cells from a human breast cancer cell line (MCF7) were treated with different compounds at several concentrations, a high-throughput microscope was used to acquire images of all treatment conditions, and these images were analyzed to extract features from individual cells after segmenting them. Segmentation and feature extraction were performed using CellProfiler. Several categories of features were extracted, including size, intensity, shape, texture and neighborhood information. More than 400 features in total were extracted per cell. The simplest algorithm comprised averaging these single cell measurements across all cells for each each treatment condition (a compound at a given concentration) to generate the treatment ‘profile’.

These profiles were evaluated on the task of classifying each treatment condition into its ‘mechanism-of-action’ (MOA) - a label that was assigned to each compound based on prior knowledge. A 1-nearest neighbor classifier was used to assign each treatment to an MOA. When finding the nearest neighbor, other treatment conditions of the same compound, but different concentrations, were left out (‘leave-one-compound-out’ cross-validation).

Ljosa et al. [9] found that reducing dimensionality using factor analysis (prior to computing averages) was the best performing profiling method, yielding a mean accuracy of 94%. Singh et al. [13] followed a similar approach, and focusing on the improvement gained by correcting illumination bias in the images, showed that 90% accuracy could be achieved by directly computing averages, after the images had been corrected for the bias. Furthermore, they found that these ‘mean profiles’ seem to be more robust than ‘factor analysis profiles’.

Deep learning has been applied to profiling for profile aggregation [16], MOA classification from CellProfiler features [5] and MOA classification from raw images [6]. Zamparo et al. [16] showed that autoencoders can be used for dimensionality reduction to improve the quality of profiles. Kandaswamy et al. [5] found that classifiers trained on CellProfiler features extracted for one set of compounds learn features that can be transferred to another set of compounds. Kraus et al. [6] evaluated their method on the same BBBC021 image set as used in this paper. Using deep learning on raw images they outperformed prior classical results. However, only a small subset of the BBBC021 image set was split into single training and test sets. We hypothesize that, this smaller dataset as well as the possibility of matching to the same compound leads to overfitting.

## 3 Experimental Setup

This work used the BBBC021 [2] image set, which was the basis for the comparisons presented in Ljosa et al. [9], and is available from the Broad Bioimage Benchmark Collection [10]. We hypothesized that generic neural networks pre-trained on natural images are able to extract biologically meaningful features from microscopy images without segmenting individual cells^3^.

We tested this hypothesis with both full images and cropped images. Illumination correction was performed to reduce the effect of uneven brightness in the image ([13]). The full images were downsampled to fit the input shape of the pre-trained networks. The images have three channels corresponding to DNA, actin and tubulin stains, which do not map to RGB channels in natural images. We dealt with this using two approaches: (1) using an arbitrary mapping (DNA → R, tubulin → G and actin → B), or (2) using the network on each greyscale stain image separately and concatenating all corresponding features to build one feature vector.

We extracted features from the images by passing them through pre-trained neural networks without fine-tuning. The networks were modified by cutting off the final classification layer, so that the penultimate layer represents the feature embedding. The features were extracted with pre-trained versions of Inception-v3^4^ [15], VGG16^5^ [12], and ResNet^6^ [4]. For comparison to the classical results, the features of each treatment were averaged over the different replicate images to generate treatment profiles. This resembles the mean profiling method, used in Singh et al. [13]. As with the classical approach, the profiles were evaluated on the task of classifying each treatment condition into its MOA using a 1-nearest neighbor classifier. To ensure that the measured performance is directly comparable, we used the same leave-one-compound-out cross-validation as Ljosa et al. [9] and Singh et al. [13].

## 4 Results

We found that ImageNet pre-trained neural networks are able to extract biologically informative features from microscopy images with accuracies ranging from 55% to 91%, compared to random chance which yields 8% (Table 1). Inception-v3 achieves the best performance, though all networks extract rich features. Furthermore, the use of different stains as individual greyscale images always performs better than an arbitrary RGB mapping. This could be due to the fact that the relations between the channels of the images are different for microscopy images and natural images. Illumination correction improves the performance of almost all network and data configurations.

**Table 1:** Results of the profiles extracted using VGG16, ResNet-101, ResNet-152 and Inception-v3. The images (1280px × 1024px) were resized to fit the input shape of each network 299px × 299px for Inception-v3, (224px × 224px for both ResNets and VGG16). Random chance would yield 8% accuracy.

|  | Inception-v3 | ResNet-152 | ResNet-101 | VGG 16 |
| --- | --- | --- | --- | --- |
| Full Images | 70.87% | 55.34% | 57.28% | 66.02% |
| + illumination corrected | 69.90% | 65.05% | 65.04% | 63.11% |
| + greyscale | 86.41% | 75.72% | 70.87% | 71.84% |
| + illumination corrected & greyscale | 91.26% | 78.64% | 79.61% | 83.50% |

The presented approach can slightly outperform the classical mean profiling method. From non-corrected images, the Inception-v3 network extracted features that achieve 86% accuracy compared to 84% using classical methods[13]. From illumination corrected images, the network generated features that achieve 91% accuracy compared to 90%. Figure 1 shows the confusion matrices for the classical and deep learning case. Interestingly, the errors of both approaches seem to be complementary, which suggests that the deep learning model captures different features than the classical method.

**Figure 1:**
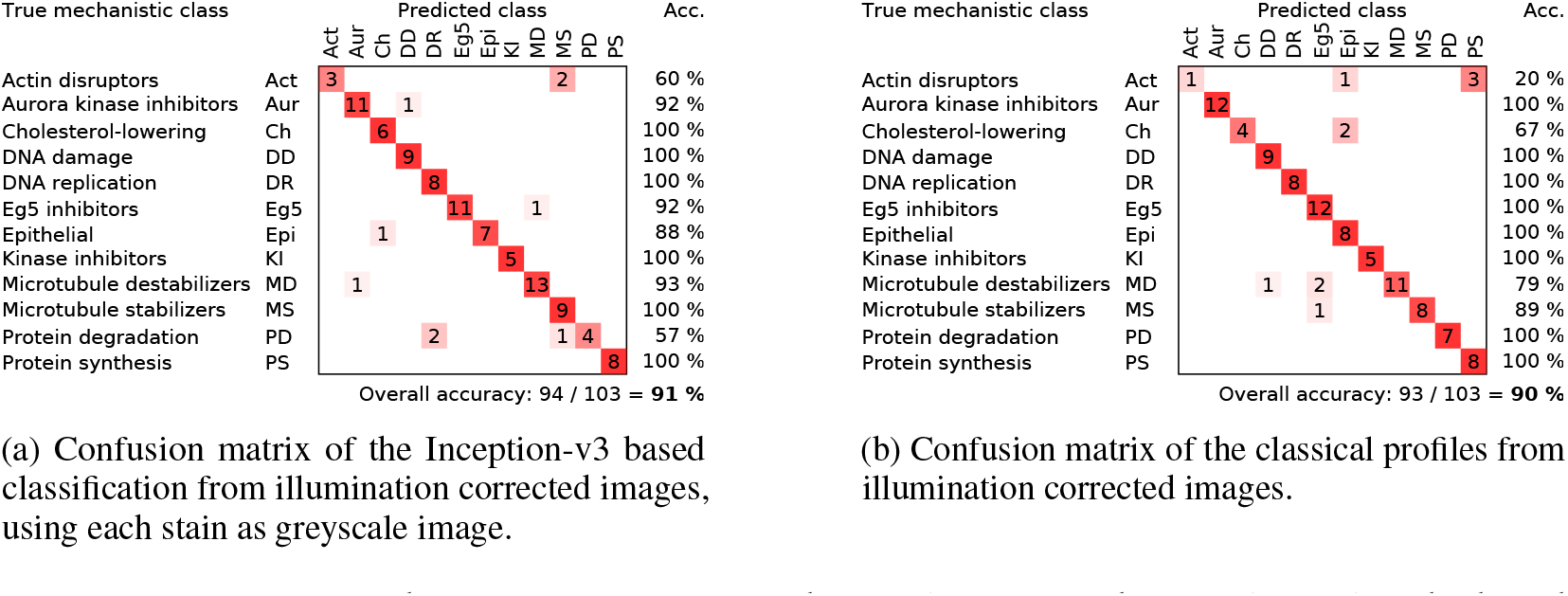
Comparison of the confusion matrices of the profiles generated using classical methods and from pretrained neural networks. The classical results were taken from Singh et al. [13]. The deep learning results were generated using the Inception-v3 architecture. Each stain was transformed as separate greyscale image.

The deep learning model also showed improvement in terms of processing time. The transformation of the images took 22 minutes using the Inception-v3 network. CellProfiler takes more than 10 hours to process the same amount of images, when running on a single core. Both algorithms were tested on a system with two 8-core Xeon CPUs, 128GB of memory and no GPU acceleration.

## 5 Conclusion

We have shown that deep networks pre-trained on natural images are capable of extracting biologically rich features from microscopy images without fine-tuning. This enables us to propose a fully automated pipeline using deep feature transfer for generating morphological profiles without human interaction. This pipeline achieves higher accuracies than previous classical methods, needs less time and expertise to extract profiles and is the first to allow for true automated high content screening by taking the human out of the loop.

We note that although the improvement in accuracy is marginal, obviating the need for an image analysis expert is a significant advantage of using deep networks. Our work builds the foundation for future explorations of transfer learning within this domain. Future work might evaluate the performance of cropped full-resolution images as well as the use of different hidden representations as extracted features. Further, fine-tuning of those feature extractors could be possible given larger microscopy image sets. We note that deep learning, usually performs better with a higher number of data points and thus hypothesize that bigger image sets will enable deep learning to flourish.

## Acknowledgements

We want to thank Mike Ando (Google, Inc.) for helpful discussions about the use of transfer learning for the domain of morphological profiling. We want to thank Allen Goodman and Claire McQuin for support with the setup of our test environment and David Dao as well as other members of the Imaging Platform for helpful discussions. AEC acknowledges NSF support (NSF CAREER DBI 1148823 to AEC).

## Footnotes

† This research has been conducted at the Broad Institute of MIT and Harvard as part of the final project of an MSc in Artificial Intelligence at the University of Edinburgh.

* Accepted as poster presentation at the 2016 NIPS Workshop on Machine Learning in Computational Biology

3 Implementation available at https://github.com/carpenterlab/2016_pawlowski_mlcb

4 Implementation from https://github.com/tensorflow/models/tree/master/inception.

5 Implementation from https://github.com/ry/tensorflow-vgg16.

6 Implementation from https://github.com/ry/tensorflow-resnet.

